# Biomimetic Virus-Like Particles to control cell functions

**DOI:** 10.1101/2024.09.14.612851

**Authors:** Hasna Maayouf, Thomas Dos Santos, Alphonse Boché, Rayane Hedna, Kaspars Tārs, Isabelle Brigaud, Tatiana Petithory, Franck Carreiras, Carole Arnold, Ambroise Lambert, Laurent Pieuchot

## Abstract

Biomimetic cues from the extracellular matrix (ECM) are essential for optimizing cell microenvironments and biomaterials. While native ECM proteins or synthetic peptides offer potential solutions, challenges such as production cost, solubility, and conformational stability limit their use. Here, we present the development of virus-like particles (VLPs) derived from the AP205 RNA phage displaying peptides from key ECM proteins and evaluate their biological activity in a variety of assays. We show that our engineered VLPs can effectively stimulate cell adhesion, migration, proliferation and differentiation. By comparing focal adhesions formed by RGD VLPs with their parent protein, fibronectin, we elucidate both similarities and differences in cell interactions. In addition, we construct heterodimeric particles co-expressing RGD with differentiation peptides and demonstrate retention of bioactivity in a multi-peptide context. This study establishes AP205 VLPs as versatile nanoscale platforms capable of tuning cell functions, with promising applications in nanomedicine and biomaterials.

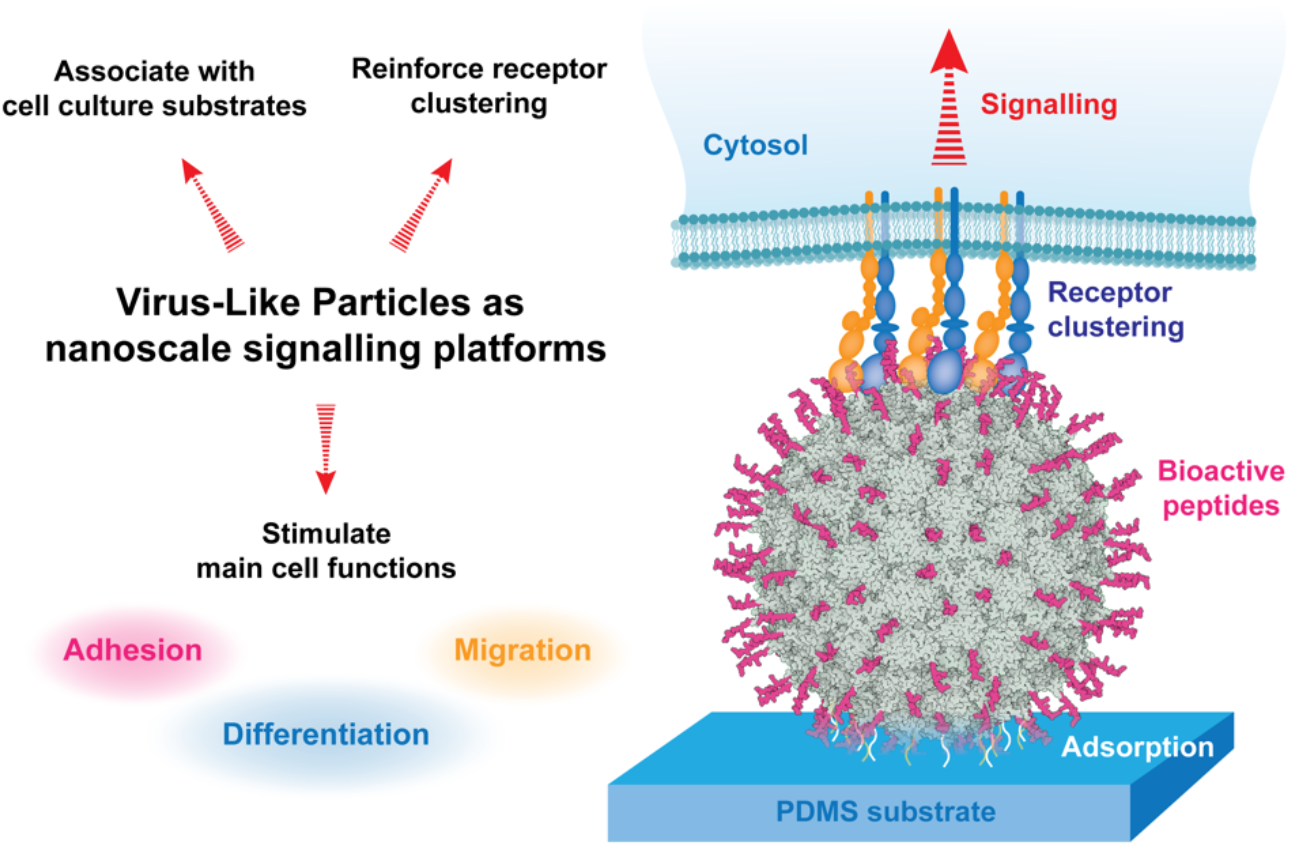
Graphical abstract.

## Introduction

In recent years, efforts have been made to develop functionalization strategies to improve the integration of biomaterials and create cellular environments that can efficiently promote cell adhesion, proliferation, and differentiation[1]. One of the main goals of this research is to mimic the *in vivo* microenvironment provided by the extracellular matrix (ECM), which regulates the physiological functions of cells[2]. Material surfaces can be functionalized with bioactive proteins from the ECM, such as collagen, fibronectin and laminin, which are known to enable cell adhesion[3]. However, the development of native protein-based systems from ECM can lead to some complications. Indeed, native proteins are often difficult to obtain (ethics, complex purification methods, high cost production), and the accessibility of their binding sites to cell receptors depends on their orientation and conformation, and the complex relationship between biochemistry, topology, and the deposition method[4]. This can be overcome by presenting short bioactive peptides derived from the full length proteins[3,5].

The bioactivity of ECM proteins comes from small domains, typically less than 50 amino acids, that are recognized as epitopes by specific cellular receptors, triggering signal transduction cascades[2]. These peptides can be synthesized or produced by recombinant technologies[6]. Their immobilization is usually achieved by adsorption or chemical grafting, for example through a covalent amide bond between their N-terminus and a carboxylic acid on the surface of the material[7].

The arginine-glycine-aspartic acid (RGD) tripeptide is one of the most studied because it promotes cell adhesion and osteointegration[8,9]. This sequence is present on many ECM proteins, including fibronectin, vitronectin and some types of collagens, which are recognized by specific receptors, the integrins[10], involved in most cell-ECM interactions. The specificity of their interaction with ECM ligands regulates their spatial distribution on cell membranes and their gathering into nanoclusters[11,12].

Nanoclustering of integrins is essential for effective signaling and depends on the global density of ECM ligands[13,14]. Integrin binding to RGD leads to the recruitment of additional integrin receptors into clusters as well as cytoskeletal components such as focal adhesion kinases (FAK), talin and paxillin, which stimulate actin polymerization[15]. Increasing RGD ligand density leads to larger adhesions and contributes to adhesion strength as well as outside-in signaling[15]. Similarly, the distribution pattern of RGD ligands directly influences cell adhesion via the constitution of these clusters[16]. Reducing the spacing of randomly arranged RGD ligands below 70 nm results in the formation of integrin clusters and enhances cell adhesion, migration and growth[16,17].

Other bioactive peptide sequences such as IKVAV and YIGSR, derived from α1 and β1 laminin chains respectively, are used to functionalize the surface of materials (e.g. implants, hydrogels, matrices) for cell adhesion, proliferation and differentiation[18–20]. Laminins are a major component of basement membranes composed of α, β and γ chains. These chains form a cross-like structure that can bind to other ECM proteins and cell receptors. Depending on the nature and combination of the three polypeptide chains, there are fifteen laminin isoforms that play an important role in maintaining the tissue phenotype^19^.

The combination of laminin with other ECM proteins such as fibronectin have been shown to modulate cell functions. For example, associating fibronectin with laminin has synergistic effects on cell differentiation[21–23]. Interestingly, this synergy is maintained when the corresponding peptides, RGD and YIGSR, are combined, further stimulating, for example, cell adhesion and proliferation of HUVECs[24,25]. These two peptides also appear to increase cell attachment, proliferation and differentiation of C2C12 skeletal myoblast cells cultured on flat and grooved PLGA films[26].

Many cellular functions are regulated by the interaction between the integrins and growth factor (GF) receptors. Integrins can directly associate with GF receptors to promote their activation and orchestrate their endocytosis and recycling[27,28]. Activation of integrin and GF receptors elicit the same signaling pathways and mechanisms involved in physiological and pathological processes[29]. Additionally, the cross-talk between these pathways regulates cell proliferation, differentiation, and migration *in vivo*. For example, insulin and integrin αvβ3 can cooperatively upregulate IGF-1 (insulin-like growth factor-1) receptor to promote smooth muscle cell proliferation[30]. In contrast, the proliferation of renal epithelial cells stimulated by EGF (epidermal growth factor) depends on α5/β1 integrins[31]. Furthermore, the association of fibronectin and BMP2 (bone morphogenetic protein 2) has been shown to have a synergistic effect on the osteogenic differentiation of C2C12 cells on hydroxyapatite coatings[32]. Additionally, RGD and BMP2 peptides derived from these proteins enhance the osteogenic commitment of human bone marrow stem cells (hBMSCs)[33]. Therefore, it is important to develop interplay systems to simultaneously present different peptides for effective bioactivity and cell signaling.

In addition to the interplay between GFs and integrin receptors, ECM proteins also provide protection and release of several matrix-bound GFs, including BMPs, which are members of the transforming growth factor-beta (TGF-β) superfamily. By binding to GFs, ECM proteins regulate their accessibility and activity until they are released by enzymatic processes[34]. For example, the cleavage of BMP2 by extracellular matrix metalloproteinases (MMPs) ensures its activity and binding to cellular receptors, which regulate bone regeneration[34].

However, it can be difficult to control and maintain the spatial distribution or orientation of peptides on material surfaces at the nanometric scale, either by chemical or physical adsorption. Furthermore, within their native environment, bioactive peptides from ECM proteins have a complex 3D distribution at the nanometer scale, making uniform peptide presentation on 2D surfaces less relevant[35]. Well-designed molecular scaffolds can help address these challenges. Such scaffolds can be derived from the capsid proteins of various viruses, known as virus-like particles (VLPs), or created de novo through well-organized nanoscale molecular self-assemblies[36–38].

VLPs are non-infectious, self-assembling nanoparticles composed of viral structural proteins. VLPs can adopt various symmetric structures (icosahedral, spherical, helical or hybrid) that typically range in size from 20 to 200 nm[39]. The varied structures of VLPs make them useful for many applications, such as vaccines and drug delivery[40]. For example, VLPs from the AP205 RNA bacteriophage have been widely studied as platforms to display repetitive peptides on their surface. These 30 nm icosahedral structures composed of 180 subunits expose the N- and C-termini of their capsid coat protein (CP3) to the outside of the particle in a highly ordered fashion, making them promising candidates for biotechnologies development. Although these VLPs have been extensively used for vaccines[38], they have not yet been applied to biomaterials.

Here we develop a series of bioactive particles based on AP205 VLPs and demonstrate that they modulate cell adhesion, proliferation, migration kinetics, and differentiation. We design heterodimeric particles co-expressing RGD together with YIGSR or BMP2 peptides and test their bioactivity. We show that the activity of RGD can be preserved when combined with other peptides on the same particle, suggesting that AP205 VLPs are a promising backbone for developing multifunctional nanoscale signaling platforms to control multiple cell functions.

## Results

### Design and production of bioactive VLPs

We genetically fused a 6x histidine purification tag to the N-terminus of the CP3 coat protein of AP205 and the following bioactive peptides to its C-terminus: RGD from fibronectin, YIGSR from laminin or an epitope derived from BMP2 (amino acids 73 to 92, Figure 1A)[41,42]. We developed heteromeric particles that expose combinations of these peptides using a co-expression vector. A first CP3 monomer was fused to a 6x histidine tag at the N-terminus and a RGD peptide at the C-terminus. A second CP3 was fused to a human influenza hemagglutinin (HA) tag at the N-terminus and either YIGSR or the BMP2 peptide at the C-terminus (Figure 1B).

**Figure 1.**
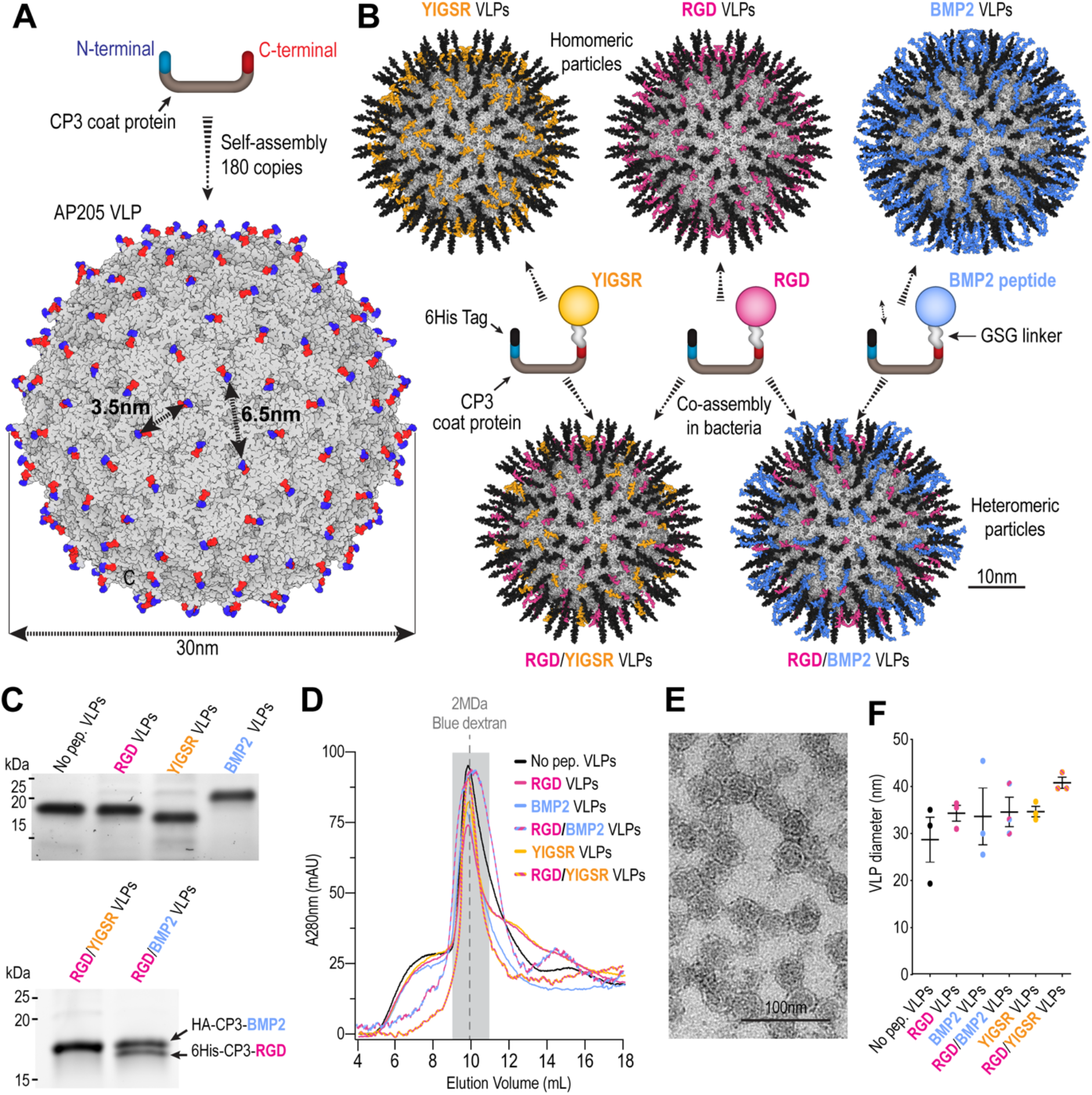
Design, production, and characterization of bioactive Virus-Like Particles. A and B: 3D renders of the original AP205 capsid (A) and the VLP variants (B). C: SDS-PAGE electrophoresis gel of protein samples from the different VLPs. D: Size-exclusion profiles of the VLP variants. E: Transmission electron microscopy image of AP205 VLPs. F: Size estimation by dynamic light scattering of the VLP variants (mean ± SEM, n=3).

We submitted the five monomer sequences to Alphafold for structure prediction, generated the particles by applying icosahedral symmetry operators and performed rendering of the VLPs using Protein Imager[43]. Predicted VLP diameters were ranging from 30 nm for AP205 without peptide to 40-43 nm for the VLP variants (Figures 1B).

We expressed the constructs in bacteria and purified the VLPs by affinity followed by size-exclusion chromatography (SEC). All constructs were well expressed, with yields around a few milligrams per liter of culture (Fig. S1A), which is typical for AP205 VLPs[44]. We assessed the purity and size of the capsids by SDS-PAGE under denaturing and reducing conditions that break apart the particles. We observed distinct protein bands, between 17 and 20 kDa, and two bands of similar intensities for RGD/BMP2 heteromeric particles, corresponding to the expected sizes of the various monomers. The SEC analysis showed a major peak eluting in the expected megadalton range (Figure 1D). We also observed the elution of lower molecular weight species, which are likely unassembled subunits. Fractions corresponding to assembled particles were collected and used for further experiments. We first estimated the size of AP205 VLPs by TEM (Figure 1E) and dynamic light scattering (Figure 1F) and obtained a particle size distribution ranging from 30 nm to 40 nm, consistent with the structural predictions.

### A simple assay to test VLP bioactivity

We developed a simple assay to test the activity of the VLPs (Figure 2A) using polydimethylsiloxane (PDMS) as a substrate. PDMS has been used extensively to study cell–substrate interactions and for biomedical applications because of its biocompatibility and physico-chemical properties[45]. However, it has poor cell adhesion properties and is usually functionalized by adsorption of matrix proteins such as fibronectin. We incubated purified particles on PDMS and evaluated their adsorption by confocal microscopy after anti-His immunostaining. We first measured the fluorescence obtained with a range of concentrations of RGD VLPs (Figure 2B). A weak coating was observed at the lowest concentration (0.65µM), with some aggregates visible on the surface. The particles then formed a more homogeneous and dense coating with increasing concentrations, with fluorescence intensity reaching a plateau at the highest concentration tested (65µM), suggesting that the PDMS binding capacity was saturated. We then compared the adsorption of the different VLP variants at 65µM and found that they associate with the surface in a similar manner (Figure S1B). Therefore, we used this concentration to functionalize the PDMS in all subsequent cell adhesion experiments.

**Figure 2.**
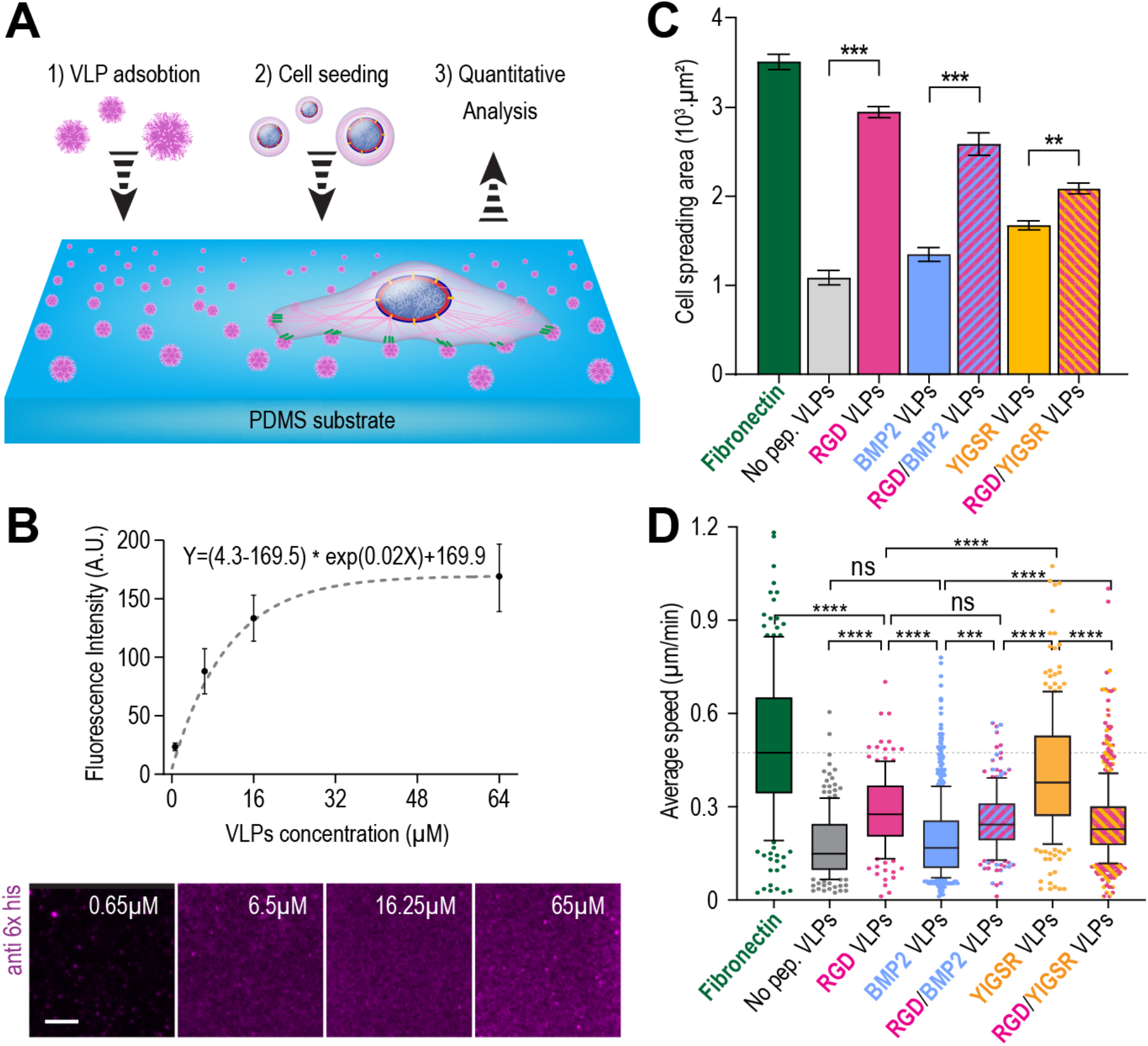
Bioactive VLPs can promote cell spreading and migration. A: Bioactivity assay based on VLP adsorption on PDMS substrates. B: Quantification of RGD VLPs adsorption on PDMS after anti-his immunostaining. Data are represented as mean ± SD with nonlinear regression fitting. Scale bar: 10 µm. C: Quantification of cell spreading on fibronectin or on the VLP variants after 18 hours of culture (mean ± SEM). (D) Average speed of C2C12 cells migrating on fibronectin and VLPs (mean ± SD, n > 140 cells). P-values were obtained using one-way ANOVA with Tukey’s multiple comparisons test with the following significances: *p < 0.05, **p < 0.01, ***p < 0.005, ****p < 0.001.

### Bioactive VLPs promote cell spreading

We compared the ability of the VLP variants to promote cell spreading using mouse C2C12 myoblasts, a cell type that can differentiate into skeletal muscle cells and osteoblasts under appropriate culture conditions (Fig. 2C). Cells were cultured for 18 hours on PDMS substrates coated with either VLP variants or fibronectin (65µM, saturating conditions). VLPs exposing RGD significantly increased cell spreading compared to control VLPs (no peptide), BMP2 VLPs or YIGSR VLPs, although cell area was lower than on fibronectin. Additionally, RGD VLPs promoted cell adhesion and spreading in the absence of serum, which typically contains adhesive molecules such as fibronectin (Fig. S2). This highlights the potential of RGD VLPs to promote cell adhesion without relying on exogenous adhesive molecules. Furthermore, heteromeric particles co-expressing YIGSR or BMP2 together with RGD peptides induced greater spreading than homomeric BMP2 VLPs and YIGSR VLPs (Fig. 2C), suggesting that RGD peptide activity is conserved.

### Bioactive VLPs influence cell migration

We analyzed the effect of VLP variants on cell migration. Cells were monitored by confocal microscopy after seeding on functionalized PDMS for 18 hours. Cell trajectories were quantified using the nucleus as a position marker to evaluate the speed, directional persistence, and total migration distance of each cell over time. Cells migrated actively on all substrates but with different dynamics (Figure 2D). Higher speeds were observed on fibronectin and particles presenting YIGSR peptides. In contrast, significantly lower cell velocity was observed on RGD VLPs. Quantification of directional persistence showed no difference between substrates (Fig. S4B). However, in terms of total migration distance, cells migrated more on fibronectin compared to VLP-coated surfaces (Fig. S4A, C). The presence of RGD at high density may facilitate more robust and stable focal adhesions (FAs) and therefore stronger attachment as well as a decrease in velocity during C2C12 migration. Consistently, the nuclei surface area was higher on fibronectin compared to RGD VLPs (Fig. S3B), confirming that cells spread more on these surfaces.

### RGD VLPs stimulate Integrin β1 recruitment at the cell periphery

We conducted a series of analyses to better characterize the focal adhesions formed on RGD VLPs, comparing them to those formed on fibronectin-coated substrates. Using vinculin, a protein involved in FA formation, as a marker, we analyzed the number, distribution, and intensity of FAs from the nucleus to the cell periphery (Fig. 3). Cell morphology analysis showed increased polarity and a migratory phenotype for cells on fibronectin compared to those on RGD VLPs (Fig. 3A). Cells adhering to RGD VLP-coated surfaces exhibited more FA clusters, which were predominantly localized at the periphery compared to those on fibronectin (Figures 3C, D and E). However, there were no significant differences in the total number, length, and mean intensity distribution of FAs per cell between the two conditions (Fig. 3C and S3).

**Figure 3.**
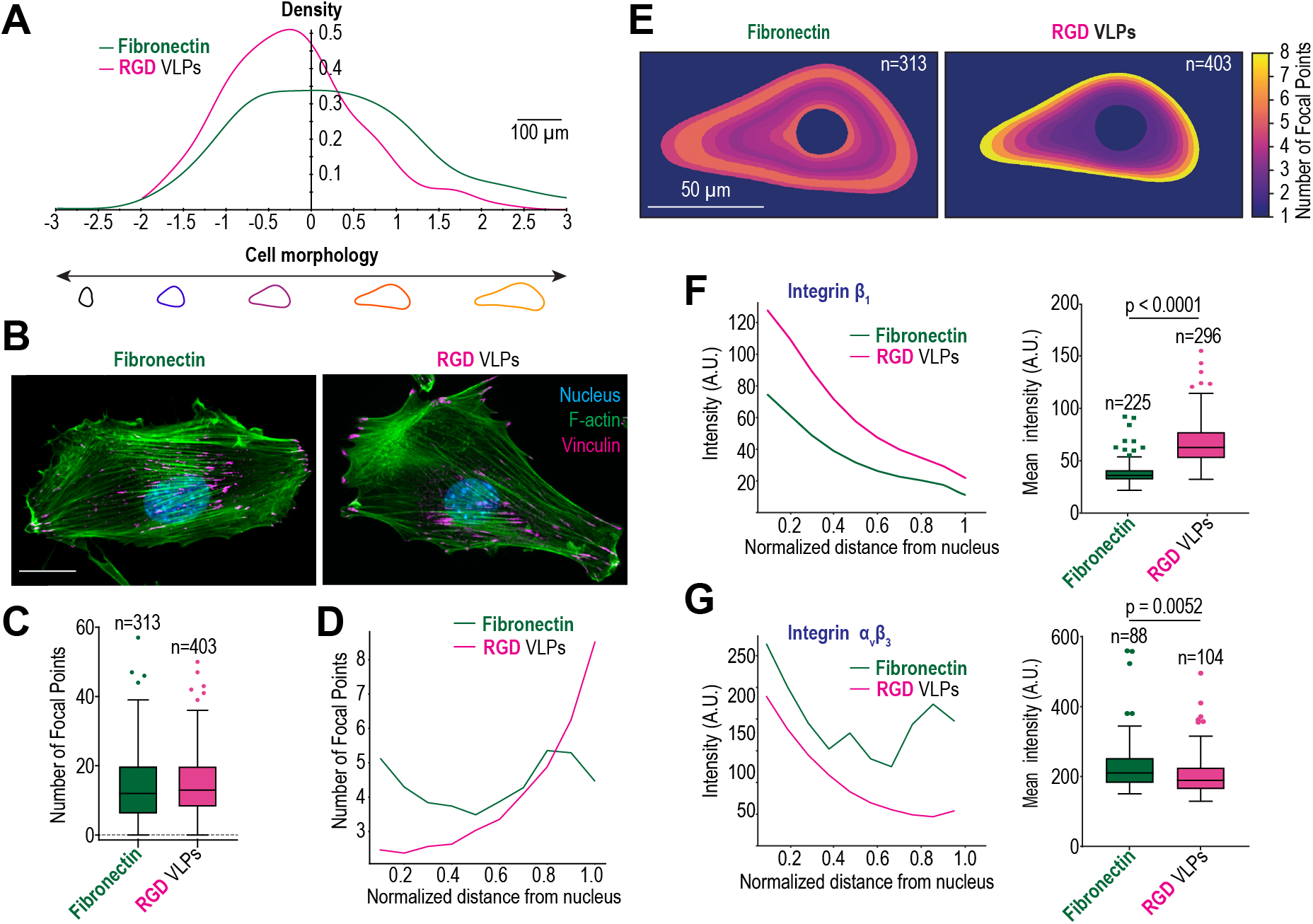
Focal adhesion density and distribution on fibronectin and RGD VLPs. A: Cell morphology distributions of C2C12 cells 18 hours after seeding on PDMS functionalized with fibronectin or RGD VLPs, B: Immunolabeling of vinculin (magenta), F-actin filaments (green), and the nucleus (blue) of C2C12 cells 18 hours after seeding on PDMS functionalized with fibronectin or RGD VLPs. Scale bar: 30µm. C: Number of focal adhesions per conditions (mean ± SD) and (D) distribution of focal adhesion number as a function of the normalized distance from the nuclear periphery (0) to the cell periphery (1). E: Cell morphology and number of focal adhesion distribution of C2C12 cells grown on fibronectin or RGD VLPs. F: Integrin β1 and (G) αvβ3 mean intensity distribution as a function of the normalized distance from the nucleus to the membrane, as well as the overall cytoplasmic mean intensity (mean ± SD). P-values were determined using the t-test.

We also found that cells seeded on RGD VLPs exhibited a stronger signal after immunostaining of integrin β1 receptors compared to fibronectin (Figure 3F). This suggests that RGD VLPs stimulate the recruitment of integrin receptors and the formation of clusters in response to high RGD densities. Our findings also indicate a higher signal after immunostaining of integrin αvβ3 receptors in cells cultured on fibronectin, particularly in the perinuclear region and the cell periphery (Figure 3G).

### VLPs can influence both proliferation and differentiation processes

We evaluated the metabolic activity of C2C12 cells at 24, 48, and 72 hours (Fig. 4A) to compare their growth on polystyrene surfaces, fibronectin-coated surfaces, and VLP-coated surfaces. While cells proliferated on all VLPs, YIGSR and BMP2 VLPs exhibited slower growth, aligning with previous cell spreading observations (Fig. 2C). Conversely, RGD VLPs and heteromeric VLPs containing RGD together with YIGSR or BMP2 demonstrated superior metabolic activity and proliferation (Fig. 4B), suggesting that RGD is more potent than YIGSR in promoting C2C12 growth (Fig. 4A, B). These results confirm that RGD functionality is preserved in heteromeric VLPs and suggest that the interaction of integrins with high-density RGD sites might activate cell signaling pathways involved in cell proliferation.

**Figure 4.**
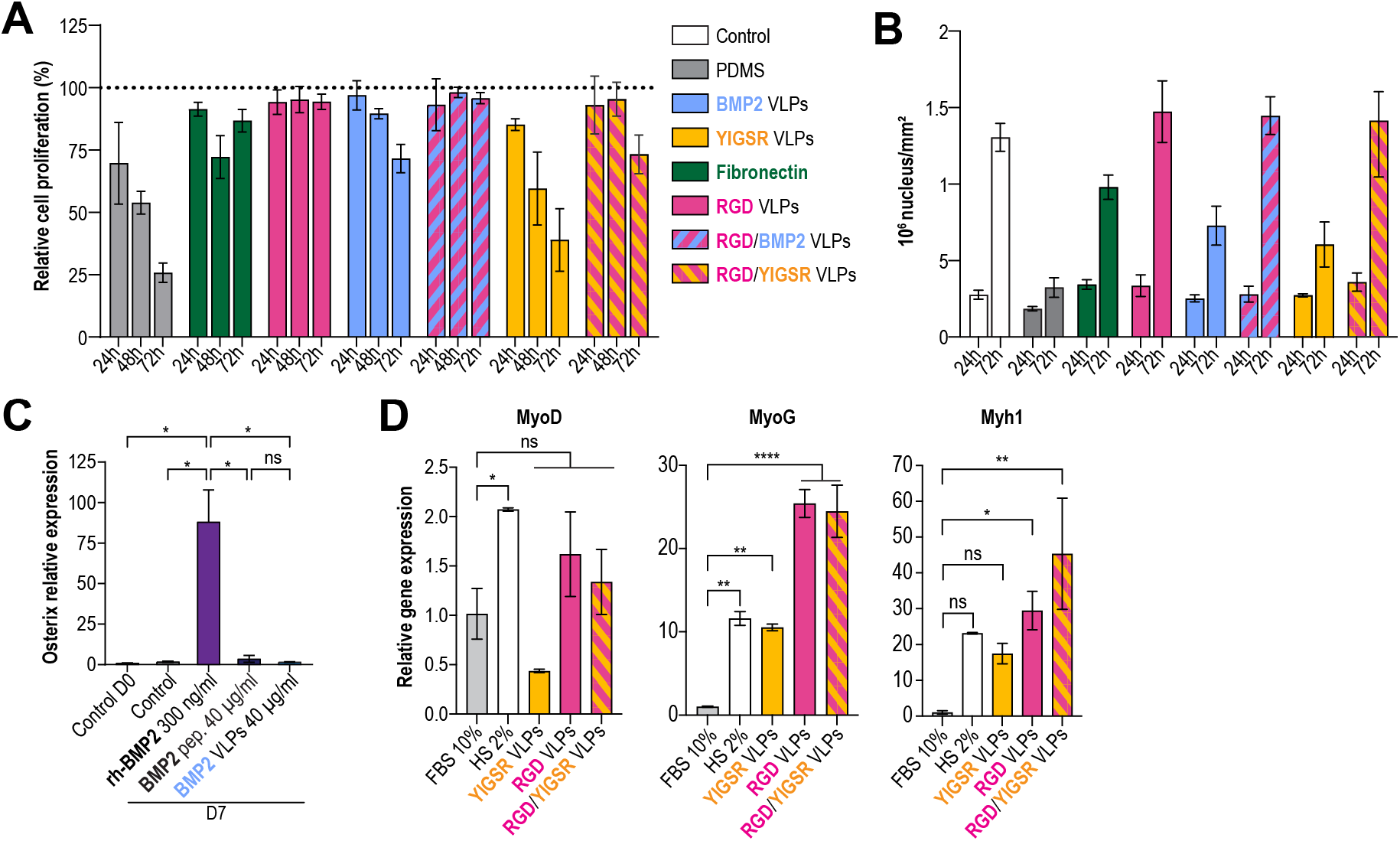
Bioactive VLPs can promote cell proliferation and differentiation. A: Relative proliferation of cells, cultured on fibronectin and on VLPs for 24, 48 and 72 hours, compared with the control without coating (mean ± SD, n = 4). B: C2C12 density (nuclei/mm^2^) on the different substrates after 24 and 72 hours (mean ± SEM, n=3). C: Effect of BMP2 VLPs on Osterix gene expression, compared to BMP2 peptide and recombinant BMP2 protein (mean ± SD). P-values were obtained using one-way ANOVA with Tukey’s multiple comparisons test (*p < 0.05). D: qPCR analysis of myogenic genes expression after 3 days of C2C12 culture in response to YIGSR, RGD and RGD/YIGSR VLPs. P-values were obtained by one-way ANOVA test with Dunnett post-hoc test comparison to the negative control (FBS 10%): *p<0.05, **p<0.01 and ***p<0.005.

To assess VLP-induced differentiation, we measured Osterix expression in C2C12 cells. Recombinant BMP2 significantly upregulated Osterix, a marker of osteogenic differentiation. However, neither BMP2 peptide nor BMP2-loaded VLPs affected Osterix expression (Figure 4C). In contrast, RGD/YIGSR VLPs stimulated the expression of MyoG and Myh1, myogenic markers indicative of myotube formation (Figure 4D). These findings suggest that the combined effects of these peptides on VLPs could potentially guide C2C12 cells towards a myogenic lineage.

## Discussion

Here, we present a series of bioactive molecular scaffolds based on the virus-like particle AP205. We demonstrate that these VLPs can partially mimic the cell-extracellular matrix by promoting cell spreading, migration, proliferation, and differentiation. We further show that peptides can be combined on the VLP surface while preserving their bioactivity.

### AP205 VLPs are efficient RGD-presenting scaffolds

Our results show that RGD VLPs enhance cell spreading and motility, although less effectively than fibronectin. Given that each 30 nm VLP presents approximately 180 RGD peptides, with only few nanometers between ligands, we hypothesize that several integrins could simultaneously bind a single VLP (see figure S6 where a RGD VLP which depicts an RGD). Previous work has shown that cell spreading is reduced when the distance between integrin ligands is greater than 58 nm[46] and that enhanced adhesion is observed when the distance between integrin molecules is less than 70 nm[47]. The high density of integrin ligands on the VLP surface may induce autonomous integrin clustering. Indeed, our results support this hypothesis by showing that RGD VLPs promote the formation and accumulation of integrin β1 at the cell periphery with lower levels of αvβ3 integrin and reduced cell motility than fibronectin. This may have potential applications in cancer research, for example by limiting cancer cell migration.

### Heteromeric VLPs are a promising strategy for combining bioactive peptides at the nanoscale

ECM-mediated synergistic signaling between adhesion receptors and GFs is essential for coordinating cellular responses[28,48]. Therefore, precise control over peptide presentation is crucial for creating microenvironments that fine-tune cell biology. We demonstrate the feasibility of producing heteromeric particles presenting different peptides by co-expressing genetic fusions of the CP3 protein. In our system, the addition of RGD peptides confers adhesive and proliferative properties to both YIGSR and BMP2 particles.

### Biomimetic VLPs: an accessible approach to functionalize surfaces and hydrogels

A key advantage of our approach is its accessibility. Any laboratory with recombinant protein expression capabilities can readily synthesize bioactive VLPs and functionalize materials. VLP yields are high, and the required resources are affordable and reusable. The spherical presentation of peptides by VLPs maximizes peptide exposure for surface functionalization. Another advantage of linking peptides to VLPs is to limit their diffusion. In the case of BMP2, for example, uncontrolled protein diffusion has been found to lead to ectopic bone formation[49].

### Current system limitations and perspectives

In our current approach, peptide stoichiometry is directly linked to promoter strength, making it challenging to precisely control peptide ratios. Producing particles with more than a few different peptides is difficult, as each version requires an additional cloning step. Additionally, the length of peptides that can be fused to the particle is constrained. For instance, our BMP2 particles failed to induce osteogenic differentiation, likely due to the peptide’s inability to mimic the complex 3D structure of the full-length protein required for BMP signaling[50]. To address these limitations, we are exploring the use of conjugation systems like SpyTag-SpyCatcher to add peptides in a second step, enabling precise control over peptide stoichiometry.

Overall, our study highlights the potential of VLPs as versatile platforms for controlling cell behavior by presenting bioactive peptide nanoclusters and mimicking cell-matrix interactions. This technology holds promise for developing novel biomaterials for tissue regeneration and wound healing.

## Methods

### AP205 VLPs molecular cloning design

The sequence coding for CP3 protein (AP205 capsid) was fused by PCR to a 6-histidine purification tag in N-term and bioactive peptides (RGD, YIGSR and BMP2) through a “GSG” flexible linker in C-term and cloned into the pET15b vector. The pETDuet-1 vector was used for the co-expression of his tagged AP205-RGD and HA tagged AP205-BMP2 or AP205-YIGSR. The primers used for amplification are listed below. Sequence ofthe inserts were verified by Sanger sequencing before their expression and production in *E*.*Coli* BL21 strain.

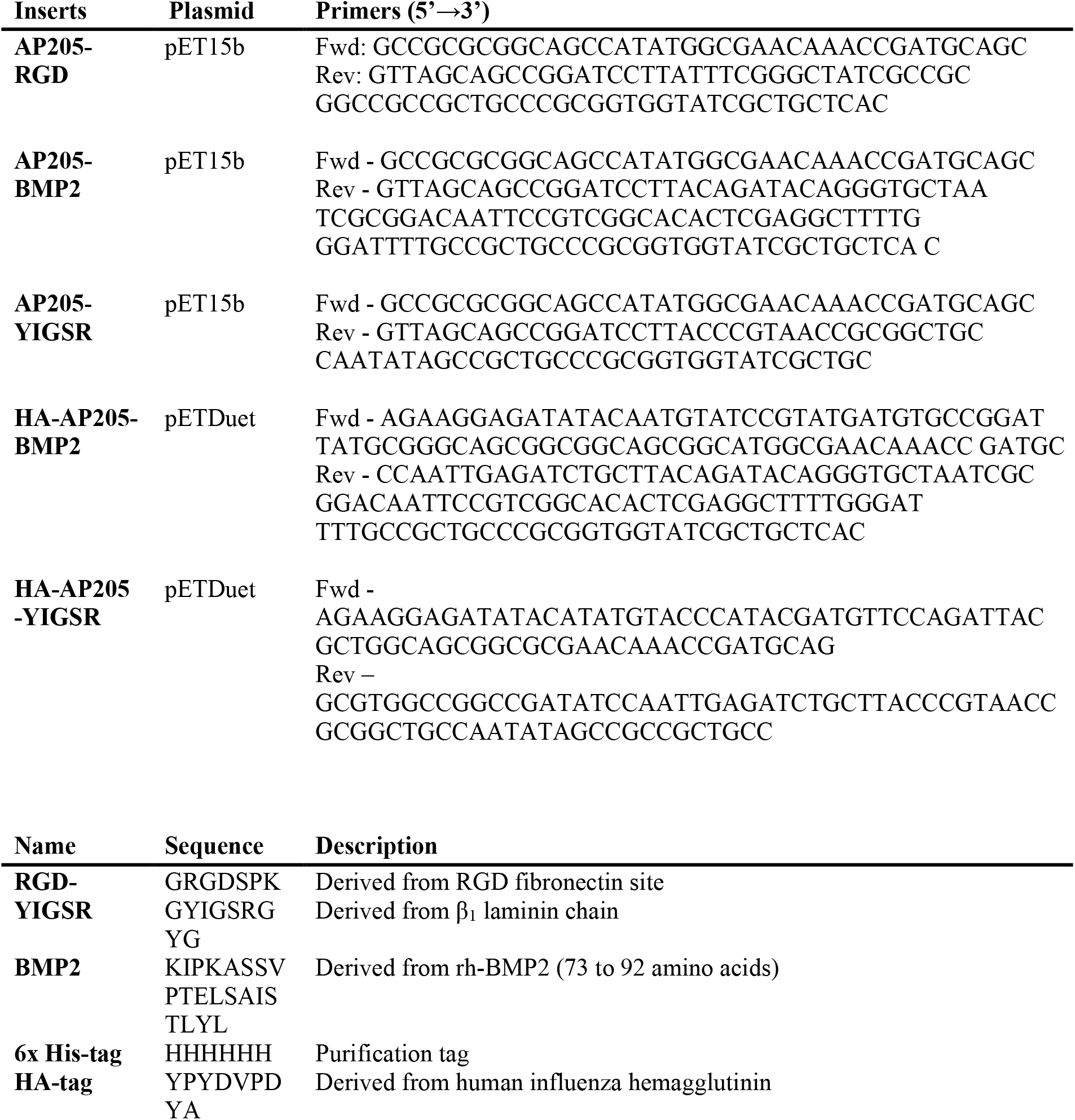

### Expression and purification of VLPs

E. coli BL21 (One Shot™ BL21 Star™) cultures were grown in Lysogeny Broth (LB) medium supplemented with ampicillin (100 µg/mL) at 37°C with shaking (160 rpm) to an OD600 of 0.6-0.8. Protein expression was induced with 1 mM isopropyl β-D-1-thiogalactopyranoside (IPTG, Sigma™) for 4 hours at 37°C in 1 L of LB medium containing 100 µg/mL ampicillin. Cells were harvested by centrifugation (4,000 x g) and stored at -20°C. After thawing on ice, cell pellets were resuspended in lysis buffer (50 mM NaH2PO4, 300 mM NaCl, 10 mM imidazole, pH 8) containing 1 mg/mL lysozyme and incubated at 4°C for 30 minutes. Cells were lysed by sonication (5 minutes, 125 watts, Q500 Sonicator, Fisherbrand™). The lysate was treated with 5 µg/mL DNase and 10 µg/mL RNase (both from Sigma™) for 15 minutes and centrifuged at 10,000 x g for 25 minutes at 4°C (JA20 rotor, Beckman Coulter). The supernatant containing the recombinant proteins was filtered through a 0.45 µm sterile filter (Whatman Puradisc). The His-tagged particles were purified by affinity chromatography on Ni-NTA agarose resin and eluted with an elution buffer containing 50 mM NaH_2_PO_4_, 300 mM NaCl, and 500 mM imidazole at pH 8. The eluted fractions were further purified by size exclusion chromatography on a 24 mL Sephacryl® 200-HR column (Sigma™) equilibrated with 1x PBS, running at a flow rate of 1 mL/min. Purified protein samples were flash-frozen in liquid nitrogen and stored at -80°C for subsequent analysis.

### Characterization of AP205 VLPs production by SDS-PAGE

20 μL of each protein sample were mixed with 20 μL of Laemmli buffer (4% w/v SDS, 20% v/v glycerol, 10% w/v beta-mercaptoethanol, 0.004% w/v bromophenol blue at pH 6.8, 125 mM Tris-HCl) and then incubated at 95°C for 10 minutes. Samples were loaded onto a Mini-PROTEAN TGX Stain-Free Precast Gel (Bio-Rad) and separated by electrophoresis. SDS-PAGE was performed at 150 V for 50 minutes. The gels were then imaged using the ChemiDoc™ imaging system (Bio-Rad). For Coomassie staining, protein samples were loaded onto Mini-PROTEAN TGX™ Precast Protein Gels (Bio-Rad). Gels were stained with Bio-Safe Coomassie Stain solution (Bio-Rad) according to the manufacturer’s recommendations.

### Quantification of VLP production by microBCA

VLP production was quantified using a Micro BCA kit (Thermo Scientific™) according to the manufacturer’s protocol For each sample and standard, 50 μL of purified protein (post-dialysis) was mixed with 150 μL of Micro BCA reagent and incubated at 37°C for 30 minutes. The absorbance of the resulting purple complex was measured at 570 nm using a Multiskan™ microplate reader (Thermo Scientific™). Protein concentrations were determined by comparison to the BSA standard curve.

### Estimation of VLP size by Dynamic Light Scattering (DLS)

Purified particles were diluted to 0.1 mg/mL in 1x PBS, filtered through a 1 µm pore size filter, and loaded into a disposable cuvette. DLS measurements were performed at a backscatter angle of 173° in triplicate at 25°C with a 60-second equilibration time using a Zetasizer Nano ZS (Malvern Panalytical). The estimated VLP diameter and polydispersity index were determined using ZS XPLORER software.

### Electron microscopy analysis

Five microliters of purified particles were adsorbed onto formvar-carbon coated grids for 10 minutes at room temperature. The grids were washed three times with pure water and stained with 2% uranyl acetate solution for 1 minute at room temperature. Excess solution was removed by blotting the grid’s edge with filter paper. Samples were then observed using a transmission electron microscope (JEOL, ARM-200F) equipped with a chemical analysis system.

### VLP adsorption on PDMS culture surfaces

PDMS was prepared by mixing the base and cross-linker (Sylgard™ 184 silicone elastomer kit) at a 10:1 ratio and pouring it into 24-well culture dishes. The dishes were cured in an oven at 80°C for 4 hours. Before use, wells were sterilized with 70% ethanol and rinsed three times with deionized water. Two hundred microliters of fibronectin or VLP suspensions at various concentrations in 1x PBS were added to PDMS surfaces and incubated for 30 minutes at room temperature. Unbound proteins were removed by washing with 1x PBS. To confirm VLP adsorption, 6xHis tags were detected using an anti-6xHis tag mouse antibody (Abcam™, 1:1000 dilution) followed by a goat anti-mouse IgG antibody conjugated to Alexa Fluor™ 650 (1:500 dilution). Fluorescence intensity was measured using confocal microscopy and quantified for each concentration substate with nonlinear regression curve fitting using the following model: Y=(*Y*_*0*_-*NS*)*exp(-*K*\**X*) + *NS*. Y0 is the binding at time zero in the units of the Y axis, NS is the binding at infinite times in the units of the Y axis and K is the rate constant in inverse units of the X axis.

### C2C12 Spreading Analysis

Mouse myoblast C2C12 cells (ATCC® CRL-1772™) were cultured in DMEM high glucose medium (Gibco™) supplemented with 10% (v/v) fetal bovine serum (FBS) and 1% (v/v) penicillin/streptomycin at 37°C with 5% CO_2_ to 70% confluence. Cells were seeded at a density of 5000 cells/cm^2^ onto fibronectin- or VLP-coated PDMS surfaces in 24-well plates and incubated for 18 hours at 37°C for cell adhesion and migration studies. All cell cultures were passaged fewer than 10 times. After 18 hours, cells were fixed with 4% paraformaldehyde for 15 minutes at room temperature. F-actin and nuclei were stained with Alexa Fluor™ 488 Phalloidin (Invitrogen™, ab150077, 1:40 dilution) and Hoechst 33342 (ThermoFisher™, 1:50 dilution), respectively, for 30 minutes at room temperature. Cells were imaged using a confocal laser scanning microscope (LSM 800 Zeiss) with 20x and 63x objectives, acquiring a series of optical sections with Zen acquisition software. Cell segmentation was performed using the Cellpose 2.0 plugin (ImageJ) with a 100 µm cell area threshold, and the cell area (µm^2^) was determined for each cell.

### Cell Imaging and Motility Assessment

C2C12 cells were labeled with SPY555 DNA fluorescent probe (Spirochrome™, 1:1000 dilution) and seeded at a density of 5000 cells/cm^2^ on PDMS substrates in complete medium. Time-lapse imaging was performed for 18 hours at 15-minute intervals using a spinning disk confocal microscope (Ti2e SD CSU W1, Nikon) with a 10x (glycerol) objective. Cell tracking was performed using Imaris® software (Oxford Instruments). Cells were detected based on fluorescence intensity with a 5 µm threshold for automatic nucleus detection. Cell position and speed were monitored over 18 hours using the autoregressive Motion algorithm.

### Proliferation and Viability Assay

C2C12 cell viability and proliferation on fibronectin- and VLP-coated PDMS were assessed using Alamar Blue (Sigma™). Cells were cultured on PDMS surfaces in 48-well plates for 24, 48, and 72 hours. The number of nuclei was counted in triplicate after 24 and 72 hours using a confocal microscope (Ti2e SD CSU W1, Nikon) with a 10x objective. Cells were fixed and stained with Alexa Fluor 488 and Hoechst before counting. The percentage of Alamar Blue reduction and relative proliferation were estimated according to manufacturer’s formulas.

### Focal Adhesions and Integrin Immunostaining

C2C12 cells were seeded at a density of 5000 cells/cm^2^ onto PDMS surfaces (24-well plate) functionalized with fibronectin or RGD VLPs (65 µM) for 18 hours. Cells were fixed with 4% paraformaldehyde for 15 minutes and permeabilized with 0.2% Triton X-100 for 7 minutes at room temperature. Non-specific binding sites were blocked with 5% bovine serum albumin (BSA) for 30 minutes at room temperature. For focal adhesion staining, cells were incubated with a primary rabbit monoclonal anti-vinculin antibody (Sigma™, 1:50 dilution) for 1 hour, followed by a secondary goat anti-rabbit Alexa Fluor™ 488 antibody (Abcam™, ab150077, 1:500 dilution). For integrin β1 staining, cells were labeled with a primary mouse anti-integrin β1 antibody (Sigma™, 1:300 dilution) and a secondary goat anti-mouse Alexa Fluor™ 647 antibody (Abcam™, ab150115, 1:500 dilution). For integrin αvβ3 labeling, cells were incubated with a primary rabbit polyclonal anti-αvβ3 antibody (ThermoFisher™, BS-1310R, 1:50 dilution) and a secondary goat anti-rabbit Chromeo™ 546 antibody (Abcam™, ab60316, 1:50 dilution). F-actin filaments and nuclei were stained with Alexa Fluor™ 488 Phalloidin and Hoechst 33342, respectively. Images were acquired using a confocal laser scanning microscope (LSM 800 Zeiss) with 20x and 63x objectives. The fluorescence intensity of vinculin and integrin β1 in individual cells was quantified using the Cellpose plugin (ImageJ) and custom algorithm define to.

### Focal adhesions and integrin cluster quantification

Focal adhesions and integrin cluster quantification was performed using the following method: Counting was conducted in each normalized zone ranging between the nucleus and the cell membrane (ranged from 0 at the nucleus periphery to 1 at the cell membrane). Normalization and counting were carried out using FIJI (ImageJ)[51]. For the analysis of β1 and αvβ3 integrin clusters, each cell was analyzed using automatic thresholding in FIJI, followed by the ‘Analyze Particles’ plugin. Cell morphology was segmented using CellPose[53], and various parameters, including fluorescence intensity and the number of focal adhesions or integrin clusters, were analyzed using CellTool[54]. We created a normalized topography from each nucleus to its cellular membrane to extract internal molecular distribution. To do so, we started on the idea to connect with the shortest distance two separated ROIs on FIJI[55,56]. To avoid artefact due to immunostaining we excluded nucleus intensity measurement in analysis.

### Myogenic differentiation assay

C2C12 cells were cultured in 48-well plates at a density of 5000 cells/cm^2^ for 3 days at 37°C with 5% CO2. For the positive control, cells were induced to differentiate into myoblasts by incubating in high glucose DMEM supplemented with 2% horse serum (HS) after seeding in complete medium for 24 hours. The untreated control group was maintained in complete DMEM medium. To assess the myogenic response to VLPs, C2C12 cells were cultured on PDMS surfaces that were coated with YIGSR, RGD and RGD/YIGSR VLPs in complete DMEM medium supplemented with 10% fetal bovine serum (FBS) for 3 days. Total RNA was extracted from the cells using a RNeasy Mini kit (Qiagen™) and quantified by measuring absorbance at 260 nm for cDNA normalization. cDNA synthesis was performed using 0.5 µg of total RNA and 4 µl of iScript RT Supermix (Bio-Rad™) in a thermocycler: 5 minutes at 25°C, 20 minutes at 46°C, and 1 minute at 95°C. The cDNA was diluted 1:20 with RNase-free water. Quantitative PCR (qPCR) reactions were set up using iTaq Universal SYBR Green Supermix (Bio-Rad™, ref 1725124) and performed on a CFX96 Touch (Bio-Rad). Mouse-specific primer sets were used for cDNA elongation at 60°C. PCR efficiencies were determined using the LinRegPCR program (version 2015.2) and relative quantification were done based on the delta CT method. *Rpl0* was used for transcript normalization.

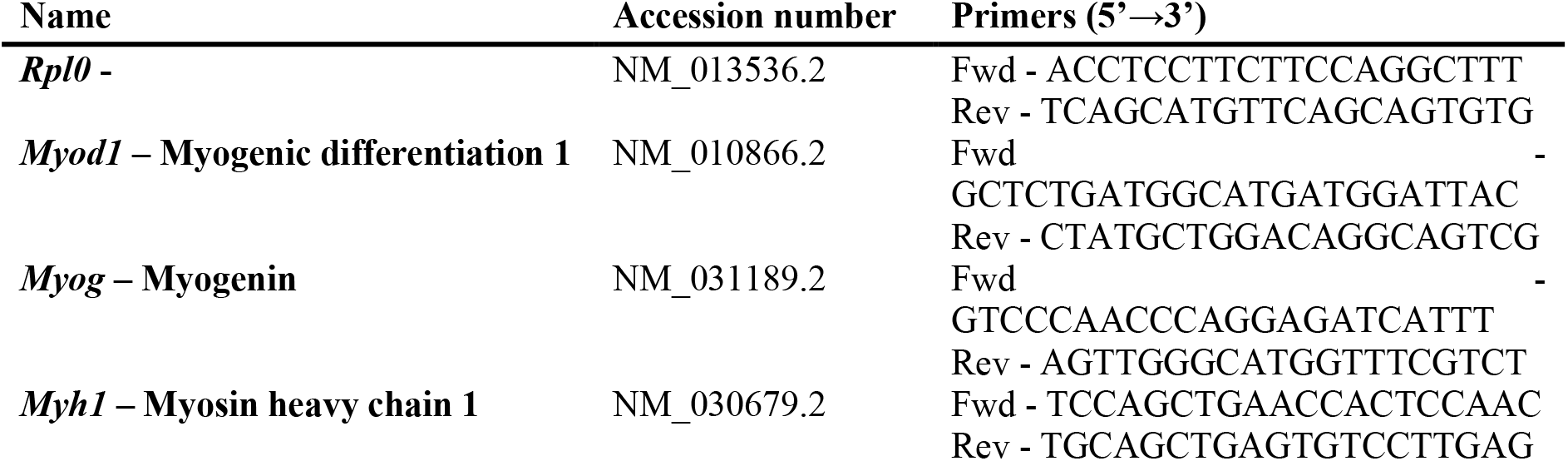

### Osteogenic differentiation assay

C2C12 cells were seeded at a density of 5000 cells/cm^2^ in 24-well plates and cultured in complete DMEM medium for 24 hours at 37°C with 5% CO2. Subsequently, the medium was replaced with either complete medium containing rh-BMP2 (Peprotech™) at 300 ng/ml, BMP2 peptides (positive control), or BMP2 VLPs at 40 µg/ml. Cells were incubated for an additional 7 days under the same conditions. Total RNA was extracted using an RNeasy Mini kit (Qiagen™). cDNA was synthesized from 0.5 µg of total RNA by RT-PCR. The synthesized cDNA was then amplified by PCR using primer sets specific for mouse osterix, rpl0, and 18S.

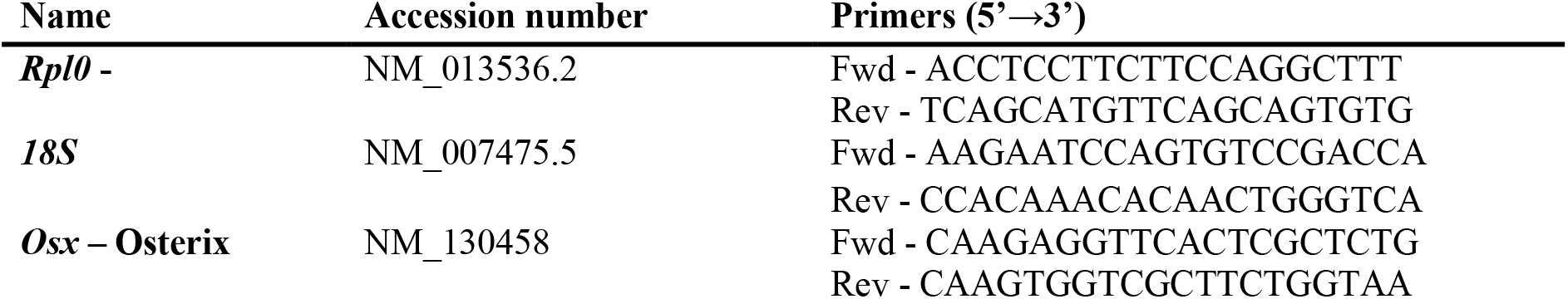

### Statistical Analysis

Each experiment was repeated at least three times. Statistical analyses were performed using t-tests, one-way and two-way ANOVA, with a significance level of 0.05 (95%). Graphs were generated using Prism software (GraphPad).

## Authors contributions

Hasna Maayouf: Experimental work and analysis, wrote the first draft of the article.

Thomas Dos Santos: Experimental work and analysis

Alphonse Boché: Focal adhesion and integrin density distribution analysis, figures plots, image analysis software design, custom code editing.

Rayane Hedna: Protein purification and gel electrophoresis.

Kaspars Tārs: VLPs structure prediction and visualization, scientific input.

Isabelle Brigaud: Gene expression analysis.

Tatiana Petithory: Imaging.

Franck Carreiras: Scientific input.

Carole Arnold: Adhesion assay on alginate hydrogel.

Ambroise Lambert: Work supervision, image analysis, manuscript editing.

Laurent Pieuchot: Conception, work supervision, wrote the final manuscript.

## Acknowledgments

We acknowledge the people running the IS2M platforms, the Agence Nationale de la Recherche (grant SPYMAT ANR-23-CE06-0026), the Centre National de la Recherche Scientifique, and the Ministère de l’Enseignement Supérieur et de la Recherche for their financial support.

## Supplementary data

**Figure S1:**
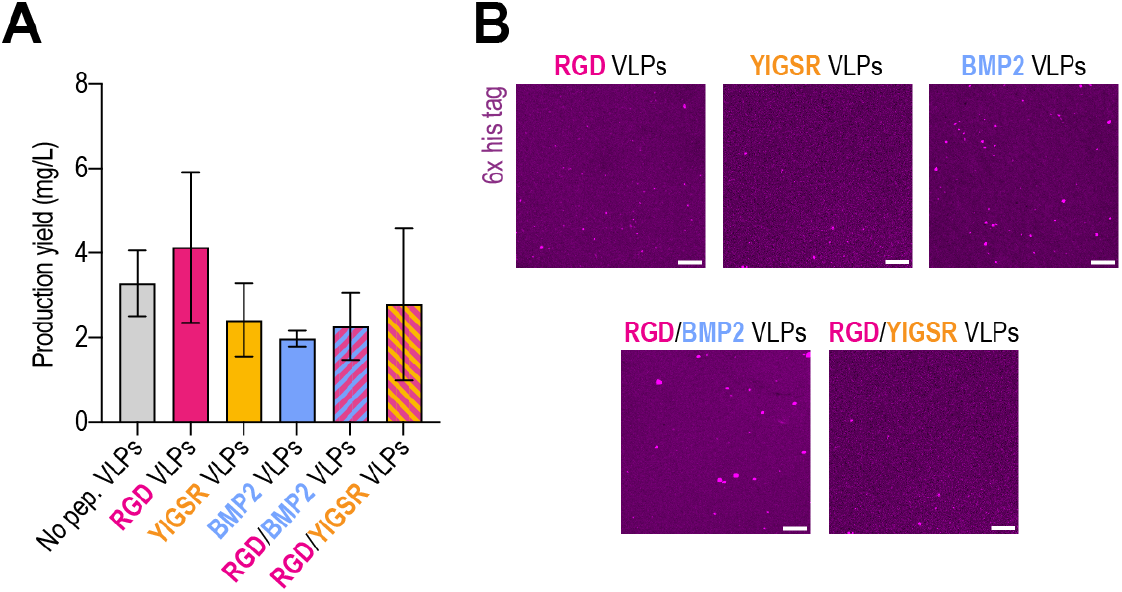
A: Total yield of purified protein per liter of bacterial culture (mg/L). Data are presented as mean ± SD (n=4). B: Immunofluorescence staining of 6x-his tags on adsorbed VLP variants on a PDMS surface.

**Figure S2:**
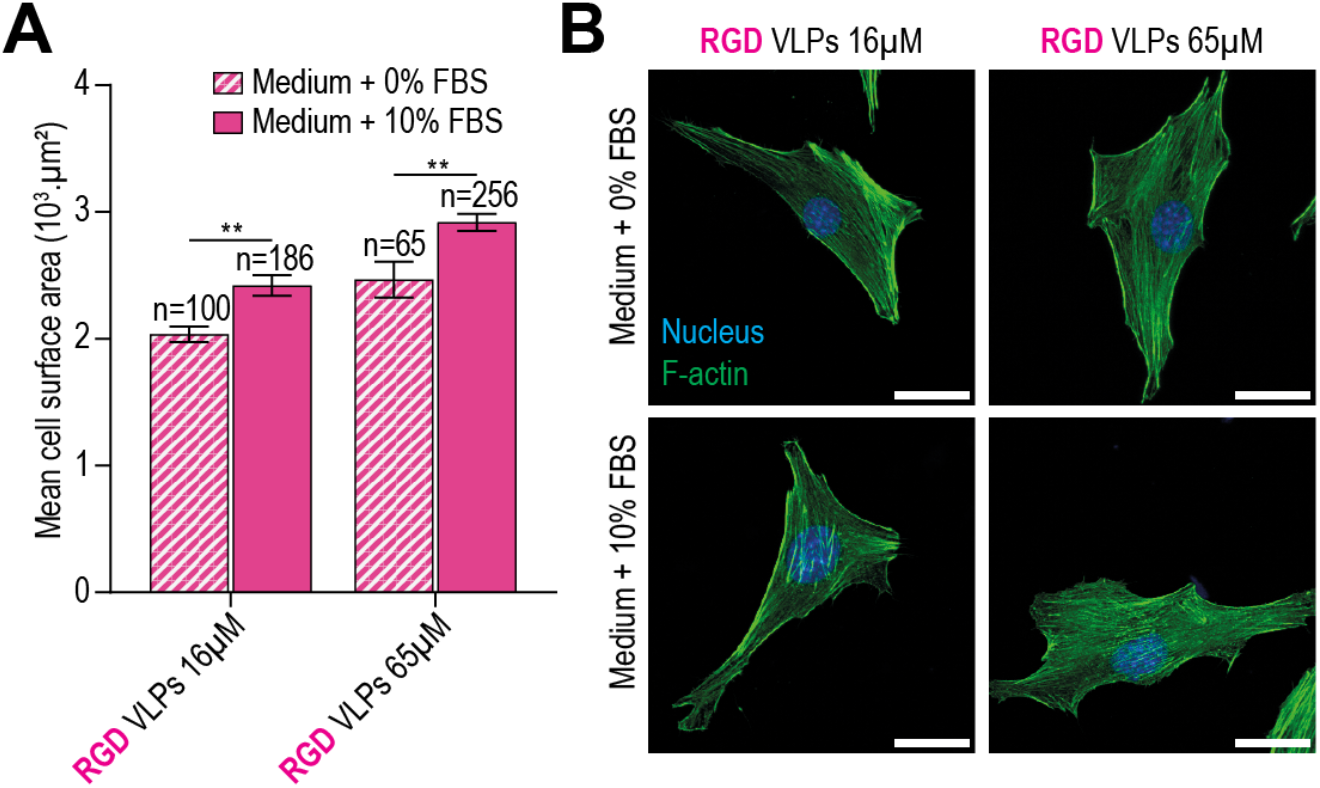
**A and B:** Quantification (A) and immunostaining (B) of C2C12 cell spreading on RGD VLPs at different concentrations after 18 hours of culture in medium supplemented with or without FBS (mean ± SEM). p-values were obtained by two-way ANOVA, ** p<0.01. Scale bar: 30µm.

**Figure S3:**
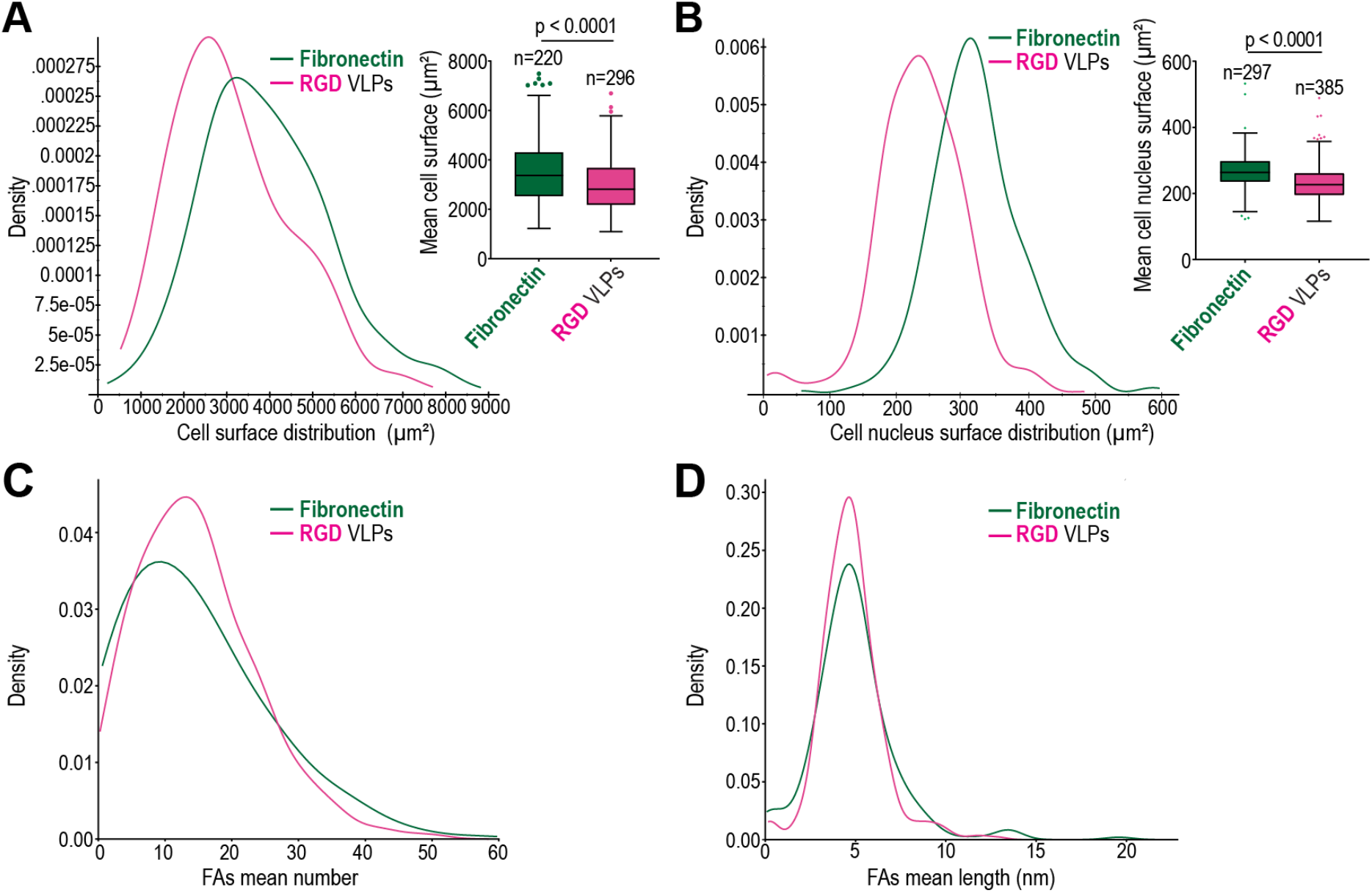
**A:** Quantification of the distribution of cell surface area and (B) nuclear area of C2C12 cells after 18 hours of culture on fibronectin or RGD VLPs analyzed using CellTool, along with mean surface area (mean ± SD, represented as box plot with Tukey whiskers). P-values were determined using the t-test. C: Total average number and (D) length of focal adhesions per cell in response to fibronectin or RGD VLPs.

**Figure S4:**
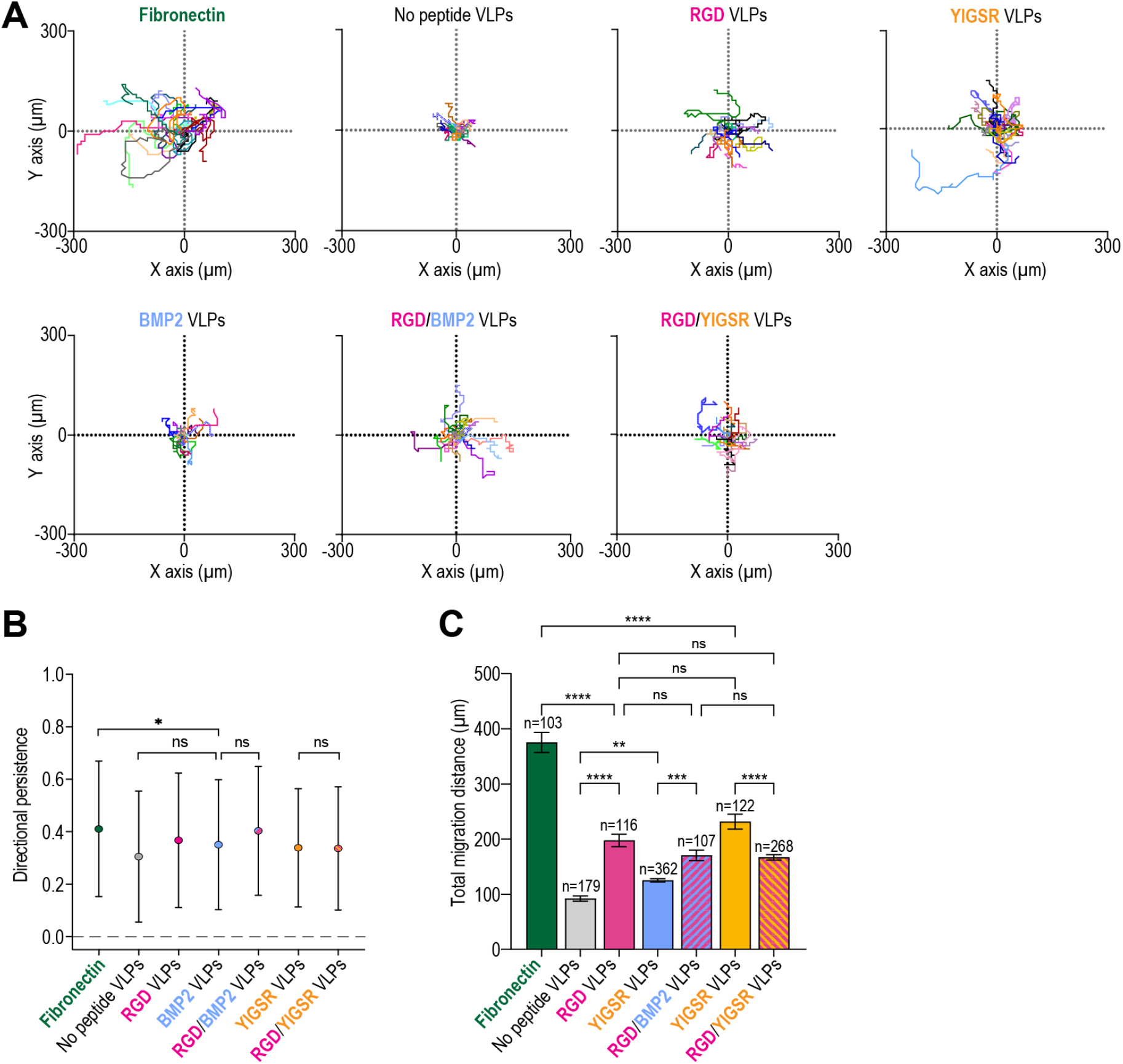
(A) Migration tracks of cells after 18 hours on fibronectin and VLP variants (n=20 in 3 independent experiments). (B) Persistence of cell migration (mean ± SD) and (C) total migration distance on fibronectin and VLP variants after 18 hours (mean ± SEM). P-values were obtained using one-way ANOVA with Tukey’s multiple comparisons test: *p < 0.05, **p < 0.01, ***p < 0.005, ****p < 0.001 (B, C).

**Figure S5:**
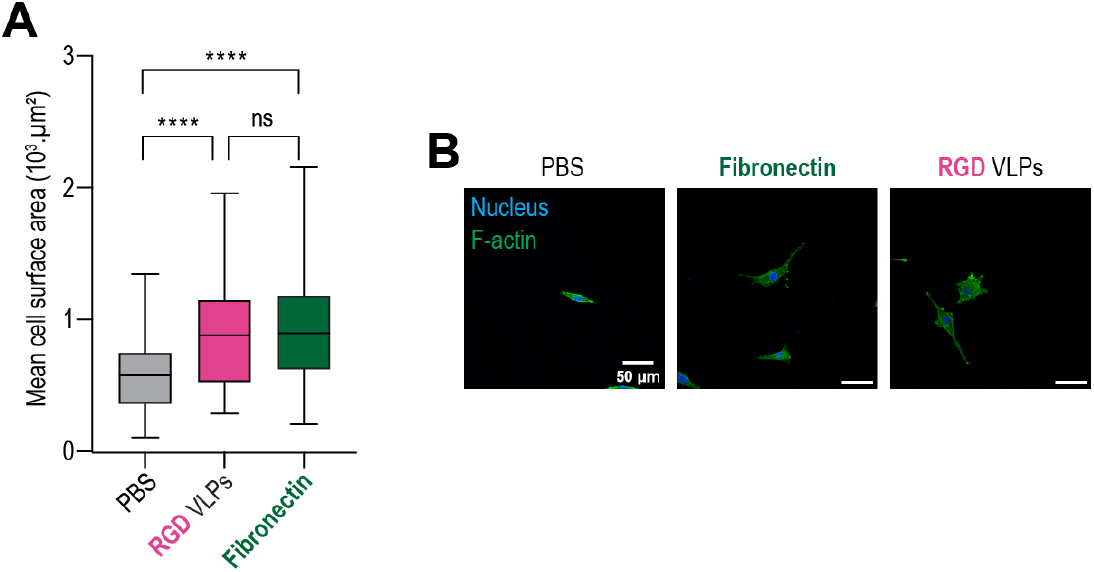
(A) Quantification of the average surface area of C2C12 cells cultured for 24 hours on 2% alginate hydrogels functionalized with PBS, RGD VLPs, or fibronectin (mean ± SD, represented as box plot). P-values were obtained by one-way ANOVA test with Tukey’s multiple comparisons test. (B) Immunofluorescence staining of C2C12 cells after 24 hours of culture on hydrogels (green: F-actin, blue: nucleus).

**Figure S6:**
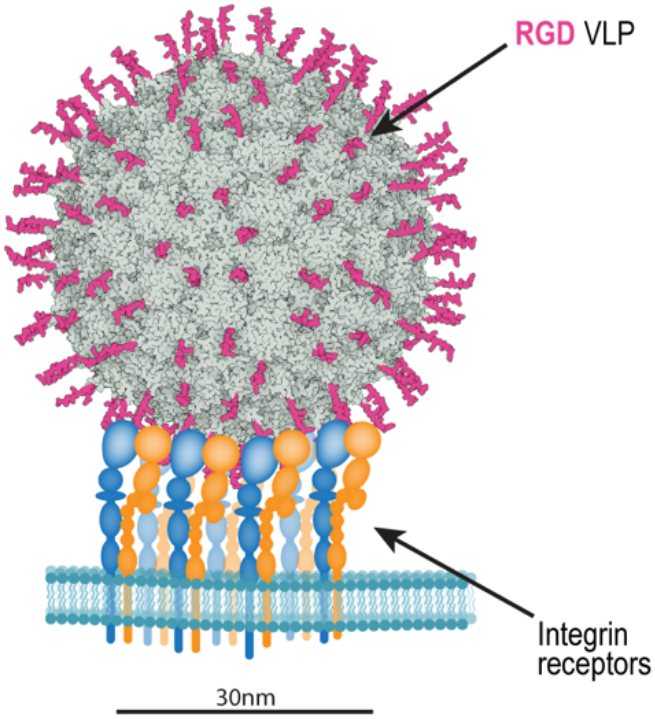
Drawing depicting an RGD VLP interacting with integrin receptors at the cell surface.

